## Appendix for "Preserved executive control in ageing: The role of literacy experience": Appendix.docx

Model 1. Random and fixed effects of the LME model with mixing cost on accuracy.

| **AIC**  -303.7 | | **BIC**  -240.9 | **logLik**  167.9 | **Deviance**  -335.7 | | | **Df. Residual**  359 | |
| --- | --- | --- | --- | --- | --- | --- | --- | --- |
| *Random effect* | | **Variable** | **Variance** | ***SD*** | | | **Corr** | |
| **Participant** | | (Intercept)  Local | 0.021  0.012 | 0.14  0.11 | | | -0.85 | |
| Residual | |  | 0.017 | 0.13 | | |  | |
| *Fixed effects****** | | ***β*** | ***SE*** | | ***df*** | ***t*** | | ***p*** |
| **Intercept** | | 0.17 | 0.05 | | 78.94 | 3.25 | | **.002 |
| **Young** | | -0.09 | 0.05 | | 90.38 | -1.94 | | .056 |
| **Local** | | -0.15 | 0.05 | | 85.43 | -2.95 | | **.004 |
| **Incongruent** | | 0.21 | 0.03 | | 249.00 | 7.29 | | ***.001 |
| **Neutral** | | 0.17 | 0.03 | | 249.00 | 5.85 | | ***.001 |
| **Print exposure** | | -0.001 | 0.002 | | 62.24 | -2.64 | | *.010 |
| **Young:Local** | | 0.09 | 0.04 | | 61.43 | 2.18 | | *.032 |
| **Young:Incongruent** | | -089 | 0.03 | | 248.07 | -2.70 | | **.007 |
| **Young:Neutral** | | -0.07 | 0.03 | | 248.08 | -2.22 | | *.027 |
| **Local:Incongruent** | | -0.11 | 0.03 | | 248.12 | -3.45 | | ***.001 |
| **Local:Neutral** | | -0.10 | 0.03 | | 248.12 | -3.03 | | **.003 |
| **Local:Print exposure** | | 0.004 | 0.002 | | 61.50 | 2.74 | | **.008 |

* Fixed effects based on effect coding, with the global condition, congruent trial and old group, coded as the intercept.

Model 2. Random and fixed effects of the LME model with mixing cost on reaction times.

| **AIC**  5548.8 | | **BIC**  5611.3 | **logLik**  -2758.4 | **Deviance**  5516.8 | | | **Df. Residual**  352 | |
| --- | --- | --- | --- | --- | --- | --- | --- | --- |
| *Random effect* | | **Variable** | **Variance** | ***SD*** | | | **Corr** | |
| **Participant** | | (Intercept)  Local | 119370  93118 | 346  305 | | | -0.29 | |
| Residual | |  | 121958 | 349 | | |  | |
| *Fixed effects****** | | ***β*** | ***SE*** | | ***df*** | ***t*** | | ***p*** |
| **Intercept** | | 1273 | 144 | | 75 | 8.86 | | ***.001 |
| **Young** | | -519 | 215 | | 62 | -2.41 | | *.019 |
| **Local** | | -470 | 153 | | 83 | -3.06 | | **.003 |
| **Incongruent** | | 103 | 65 | | 243 | 1.59 | | .114 |
| **Neutral** | | 140 | 64 | | 241 | 2.21 | | *.028 |
| **Print exposure** | | -5 | 5 | | 63 | -1.09 | | .279 |
| **Young:Local** | | 812 | 224 | | 62 | 3.62 | | ***.001 |
| **Young:Print exposure** | | 7 | 9 | | 60 | 0.79 | | .432 |
| **Local:Incongruent** | | -50 | 90 | | 241 | -0.56 | | .579 |
| **Local:Neutral** | | -264 | 89 | | 240 | -2.98 | | **.003 |
| **Local:Print exposure** | | 15 | 5 | | 64 | 3.03 | | **.004 |
| **Young:Local:Print exposure** | | -26 | 9 | | 61 | -2.74 | | **.008 |

* Fixed effects based on effect coding, with the global condition, congruent trial and old group, coded as the intercept.
